# KAS-pipe2: a flexible toolkit for exploring KAS-seq and spKAS-seq data

**DOI:** 10.1101/2022.08.10.503490

**Authors:** Ruitu Lyu, Tong Wu, Gayoung Park, Yu-Ying He, Chuan He, Mengjie Chen

**Affiliations:** Department of Chemistry, University of Chicago, IL, USA; Howard Hughes Medical Institute, University of Chicago, IL, USA; Department of Biochemistry and Molecular Biology, Institute for Biophysical Dynamics, University of Chicago, IL, USA; Department of Medicine, The University of Chicago, Chicago, IL, USA; Department of Human Genetics, The University of Chicago, Chicago, IL, USA

**Keywords:** transcription, KAS-seq, spKAS-seq, enhancers, R-loops

## Abstract

Kethoxal-assisted ssDNA sequencing (KAS-seq) is gaining popularity as a robust and effective approach to study the dynamics of transcriptionally engaged RNA polymerases through profiling of genome-wide single-stranded DNA (ssDNA). Its latest variant, spKAS-seq, a strand-specific version of KAS-seq, has been developed to map genome-wide R-loop structures by detecting imbalances of ssDNA on two strands. However, user-friendly, open-source analysis pipelines for KAS-seq data are still lacking. Here we present KAS-pipe2 as a flexible and integrated toolkit to facilitate the analysis and interpretation of KAS-seq data. KAS-pipe2 can perform standard analyses such as quality control, read alignment, and differential RNA polymerase activity analysis. In addition, KAS-pipe2 introduces many novel features, including, but not limited to: calculation of transcriptional indexes, identification of single-stranded transcribing enhancers, and high-resolution mapping of R-loops. We use benchmark datasets to demonstrate that KAS-pipe2 provides a powerful framework to study transient transcriptional regulatory programs. KAS-pipe2 is available at https://github.com/Ruitulyu/KAS-pipe2.

## Introduction

Transcription, the process of synthesizing RNA molecules from a DNA template, is mediated by three different RNA polymerases (Pols)[1, 2]. In eukaryotes, RNA Pol I and Pol III synthesize ribosomal RNAs (rRNAs) and transfer RNAs (tRNAs), respectively, whereas RNA Pol II mainly produces nascent messenger RNAs (mRNAs) of protein coding genes, which are subsequently processed into mature mRNAs and transported into the cytoplasm[3, 4]. Although transcriptional activities and their regulation can be traced indirectly by measuring steady-state RNA levels, only direct, real-time measurements of RNA polymerase or nascent RNA activity can reveal how transcription is dynamically regulated in response to different conditions. In recent years, many complementary approaches have been developed to detect nascent RNAs or to directly analyze the occupancy of RNA Pols, including 4SU RNA-seq, GRO-seq, PRO-seq, NET-seq, mNET-seq, and START-seq[5-10]. Although these approaches use different techniques, they share many of the same limitations; for example, they require millions of input cells, their ability to detect low-abundant RNA is limited, and they are susceptible to post-transcriptional RNA processing. To overcome these issues, we have developed kethoxal-assisted ssDNA sequencing (KAS-seq), a technology that can sensitively detect ‘transcription bubble’, the short stretches of unwound DNA that form when RNA Pols engage DNA for transcription. We have shown that the transient ssDNA present in transcription bubbles provides a more direct readout of transcriptional activity *in situ* than the nascent RNAs themselves[11, 12]. With the rapid (within 5 min), sensitive, and specific chemical reactions between N_3_-kethoxal and guanine bases in ssDNA, KAS-seq can work with as few as 1,000 cells or even with frozen tissues (Fig. 1a-b). We have also developed a strand-specific version of KAS-seq (spKAS-seq), which enables the high-resolution mapping of R-loops by detecting asymmetric ssDNA exposure in the two strands.

**Figure 1.**
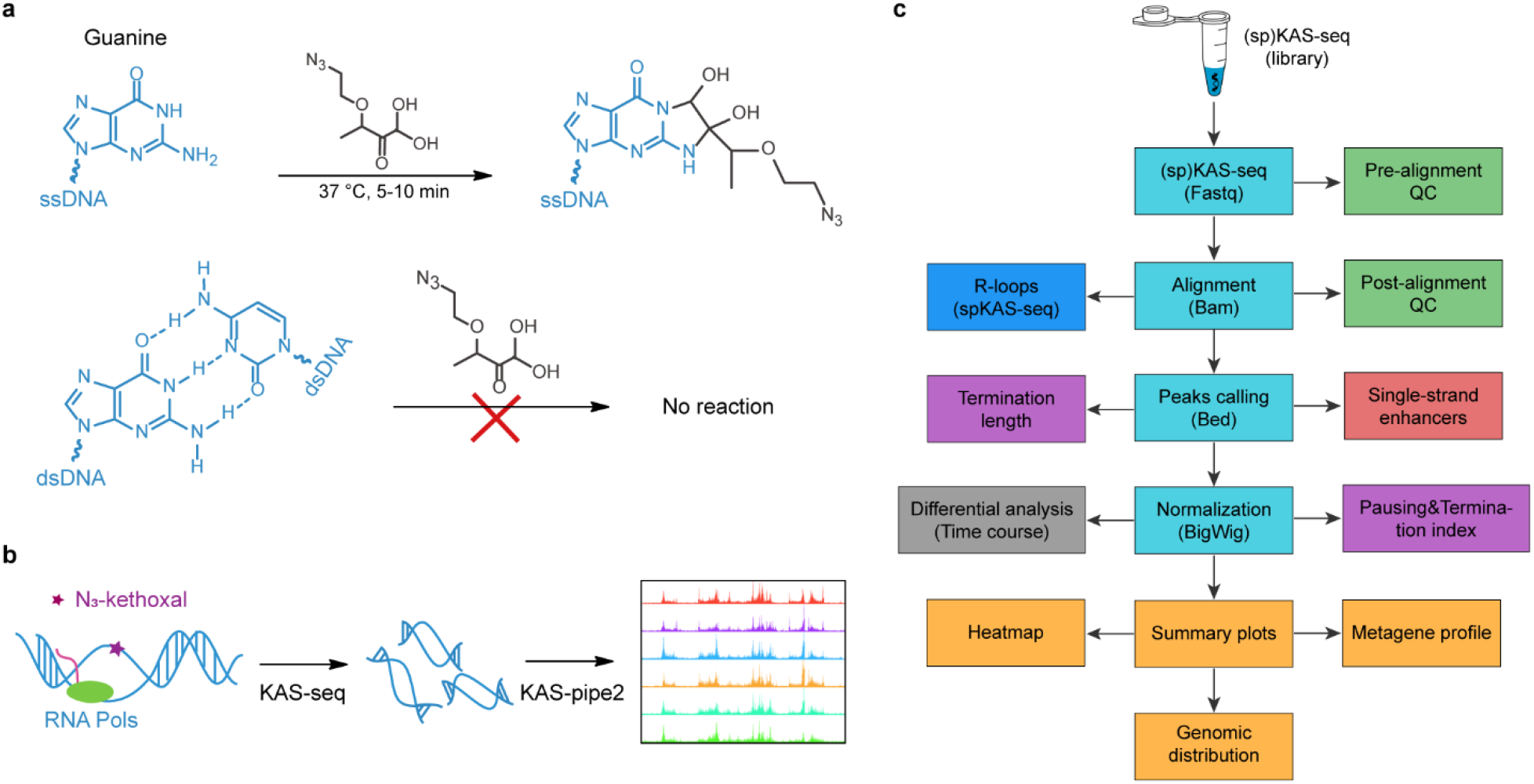
Schematic representation of KAS-seq experiments and KAS-pipe2 workflow. **a** The selectivity of N_3_-kethoxal for the guanine bases in ssDNA. Top: The fast chemical reaction kinetics between N_3_-kethoxal and exposed amine groups of guanine base. Bottom: The hydrogen bonding interactions of the Watson-Crick base pairing in double-stranded DNA (dsDNA) block the reaction between guanine and N_3_-kethoxal. **b** A schematic diagram of KAS-seq as used to map genome-wide ssDNA profiles. Purple star represents the N_3_-kethoxal labeling. Green circle represents RNA polymerases. **c** The overview of KAS-pipe2 workflow. These colored squares represent different analytical modules in the KAS-pipe2 pipeline.

Although the number of studies using KAS-seq has increased dramatically since its debut, existing bioinformatic methods are severely inadequate for processing KAS-seq data. Most relevant bioinformatics tools were developed for chromatin immunoprecipitation sequencing (ChIP-Seq) or chromatin accessibility (ATAC-seq, DNase-seq and FAIRE-seq) data[13-15]. Therefore, they cannot capture the characteristics of ssDNA profiles and will return suboptimal results when applied to KAS-seq. New methods specifically tailored to KAS-seq are needed to fully harness its power to survey the dynamics of transcriptionally engaged RNA Pols.

Here we present KAS-pipe2[16], a user-friendly, flexible and integrated toolkit for KAS-seq and spKAS-seq data analysis. The KAS-pipe2 framework combines state-of-the-art read-alignment and quantification tools with newly developed functionalities to facilitate processing and interpretation of KAS-seq data for different research purposes. We use KAS-seq data generated in 6 human cell lines (A375, HCT116, HEK293T, HeLa-S3, HepG2 and NHEK) as a benchmark dataset to demonstrate that KAS-pipe2 offers a comprehensive workflow to investigate the dynamics of transcription regulation.

## Results

### Overview of KAS-pipe2 workflow

KAS-pipe2 consists of eight functional modules, which can be used independently or collectively (Fig. 1c). With single-or paired-end raw sequencing files as input, KAS-pipe2 can be used to: (1) generate pre- and post-alignment quality control metrics; (2) perform read alignment and quantification; (3) generate summary plots of KAS-seq signals in coding regions;

(4) calculate transcription pausing, elongation, and termination indexes; (5) calculate transcription termination length; (6) identify single-stranded transcribing enhancers; (7) identify R-loops genome-wide; and (8) perform differential RNA Pols activity analysis for various study designs. By taking advantage of its modularity, users can choose to execute the entire KAS-pipe2 workflow, or choose individual analytical modules as needed.

The computational methods for these tasks listed in modules 1,2,3, and 8 are relatively well established. KAS-pipe2 integrates existing state-of-the-art tools to accomplish these tasks; for example, KAS-pipe2 uses *trim-galore* for the removal of adapter and low-quality sequences[17, 18], *MACS2* for KAS-seq peak calling[19], *deeptools* for metagene profiling and heatmaps, and *DESeq2* or *ImpulseDE2* for differential RNA Pols activity analysis[20-22]. For KAS-seq read alignment, KAS-pipe2 users can select between *BWA-MEM* and *Bowtie2*[23, 24]. Although both aligners perform well in terms of short-read alignment, they have trade-offs between mapping accuracy and speed. KAS-pipe2 utilizes new methods for the tasks listed in modules 4,5,6, and 7, because no satisfactory tools have been developed up to this point.

Results from a genomics analysis pipeline can be sensitive to the choices of tools and their tuning parameters. Thus, we have generated KAS-seq data from 6 human cell lines to provide a benchmark dataset and evaluate impacts of parameter choices. Finally, the default setting for KAS-pipe2 is a best-practice pipeline with carefully determined parameters. In the following sections, we discuss the new analysis tools implemented in KAS-pipe2 and showcase their functionalities using the benchmark dataset.

### Introduction of KAS-seq data and quality control metrics

Generally, a standard KAS-seq experiment will generate two libraries, one with enriched ssDNA through the biotin–streptavidin interactions and its matched input control (fragmented gDNA). High-quality KAS-seq samples usually have significant ssDNA enrichment over input controls. KAS-pipe2 uses a fingerprint plot and fraction of reads in peaks (FRiP) to examine whether ssDNA signals are sufficiently enriched (Fig. 2a-b). As a rule of thumb, a high-quality KAS-seq dataset should have at least 50,000 KAS-seq peaks with a FRiP value greater than 25% (Fig. 2b). If multiple biological replicates are available, KAS-pipe2 can generate correlation scatterplots to measure the consistency across replicates (Fig. 2c).

**Figure 2.**
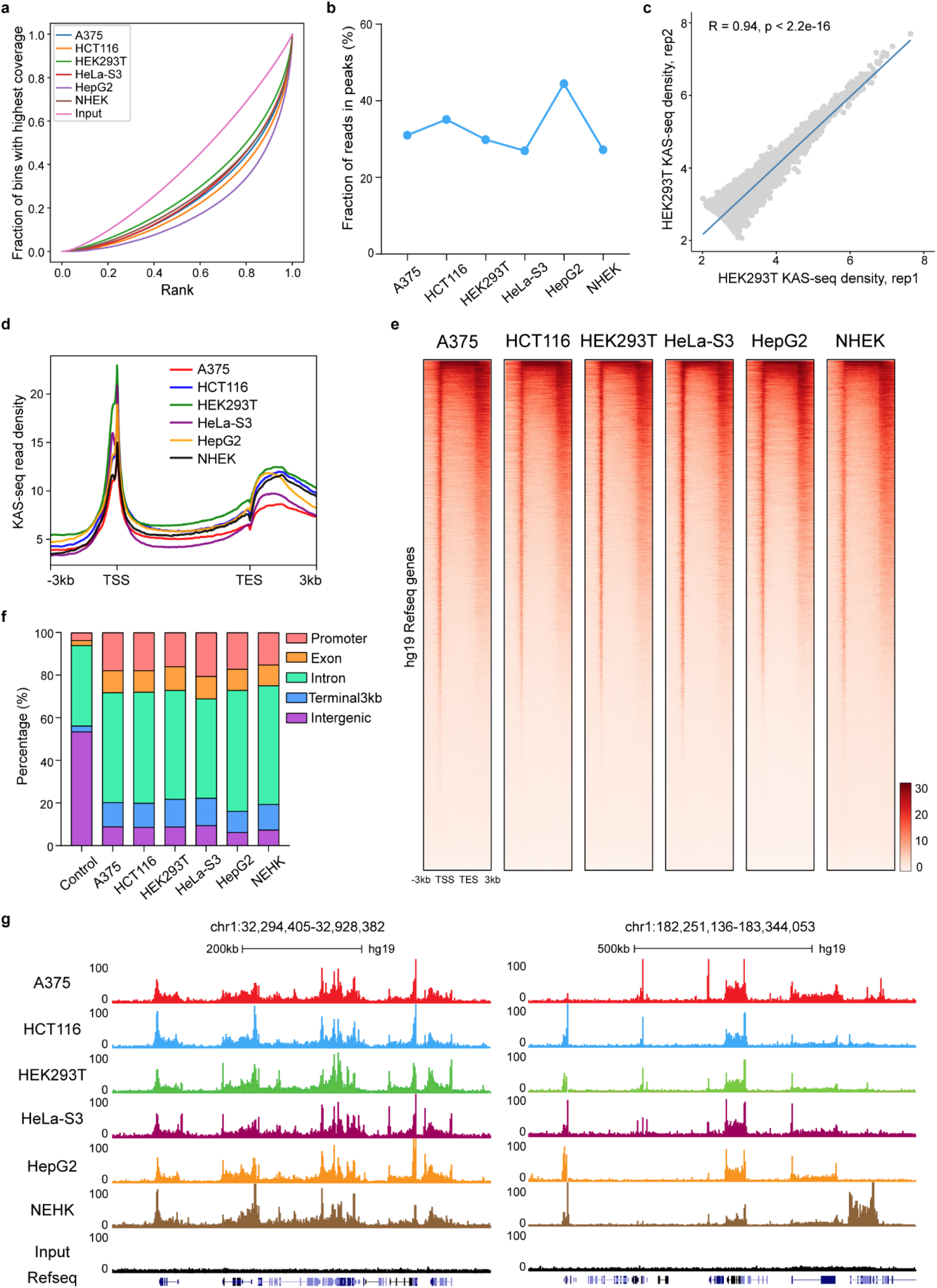
Quality control and the global view of KAS-seq data. **a** Fingerprint plot of KAS-seq data showing that KAS-seq signals can be significantly separated from the background input signals. KAS-seq data were generated in 6 benchmark human cell lines, including A375, HCT116, HEK293T, HeLa-S3, HepG2 and NHEK cell lines. **b** Fraction of reads in peaks (FRiP) metrics of KAS-seq data showing the fraction of unique KAS-seq reads mapped on KAS-seq peaks. **c** The scatterplot showing the Pearson correlation between two replicates of KAS-seq data in HEK293T cells. R value represents the Pearson correlation coefficient. P value was determined using paired samples T-test (parametric). **d-e** Metagene profile (d) and heatmap (e) plots showing the distribution of KAS-seq read density at gene-coding regions, with 3□kb upstream of TSS and 3□kb downstream of TES shown. **f** Stacked bar plot showing the percentages of KAS-seq peaks distribution on different genomic features. Promoter regions were defined as 2 kb upstream and downstream of the transcription start site (TSS). The percentage of each genomic feature in hg19 Refseq annotation is shown as the “control”. **g** Snapshots of KAS-seq data custom tracks from UCSC Genome Browser showing the pattern of ssDNA peaks on two example regions. KAS-seq data were generated in 6 benchmark human cell lines.

Previous studies show that KAS-seq exhibits a signature pattern in gene coding regions, starting with a strong and sharp peak around the transcription start site (TSS), followed by relatively moderate and broad signals that span the entire gene body, and ending with a strong but broad peak from the transcription end site to its downstream region[11]. As part of the QC procedure, KAS-pipe2 can examine this pattern by generating metagene profiles and heatmaps for KAS-seq signals in coding regions (Fig. 2d-e) and pie charts for the genomic distributions of KAS-seq peaks. In the benchmark dataset, we find that KAS-seq peaks are enriched on promoters (15-25%), gene bodies (50-70%), and transcription termination regions (10-15%), in each cell line (Fig. 2f). Finally, KAS-pipe2 can generate KAS-seq read-density files that can be viewed in the UCSC genome browser, providing users an additional option for QC through visual inspection (Fig. 2g).

### Quantifying the dynamics of transcriptionally engaged RNA Pol II

The process of transcription can be split into 3 main stages: initiation, elongation, and termination[25]. We define three metrics for protein coding genes that characterize the dynamics of transcriptionally engaged RNA Pol II: 1) pausing index (PI), the ratio of KAS-seq read density at the promoter-proximal region (TSS+/-500 bp) to that in the gene body[6, 26]; 2) elongation index (EI), the averaged KAS-seq density in both the promoter-proximal region and the gene body; and 3) termination index (TI), the ratio of KAS-seq read density in the transcription termination region to the read density in the gene body (Fig. 3a). PI is a metric originally proposed for GRO-seq data, to infer the initiation rate of paused RNA Pol II to productive elongation. EI is a newly proposed metric, which can quantitatively reflect the elongation rate of transcriptionally engaged RNA Pol II. TI is proposed to measure the efficiency of RNA Pol II “releasing” from its DNA template. To facilitate the quantification of the dynamics of transcriptionally engaged RNA Pol II, we implemented new functions in KAS-pipe2 to calculate PI, EI, and TI values for each protein-coding gene in KAS-seq data. We subsequently applied KAS-pipe2 to the benchmark dataset and summarized all three metrics for 6 human cell lines (Fig. 3b-d). Compared to other cell lines, HeLa-S3 cells have the most genes (8,771) with high PI values (no less than 2), but an average number of genes with high EI and TI values (Fig. 3b). This observation suggests that RNA Pol II in HeLa-S3 cells may be more stably paused at promoter-proximal regions and is tightly regulated by the transcription factors that control entry into productive elongation. Furthermore, we found that the number of genes with high EI values were very similar across cell lines, while TI values varied widely between cell lines (Fig. 3c-d). This finding indicates that the transcription termination process is dynamically regulated. Overall, these three metrics provide a quantitative summary of RNA Pol II activities at different stages of the transcription cycle (Fig. 3e-i).

**Figure 3.**
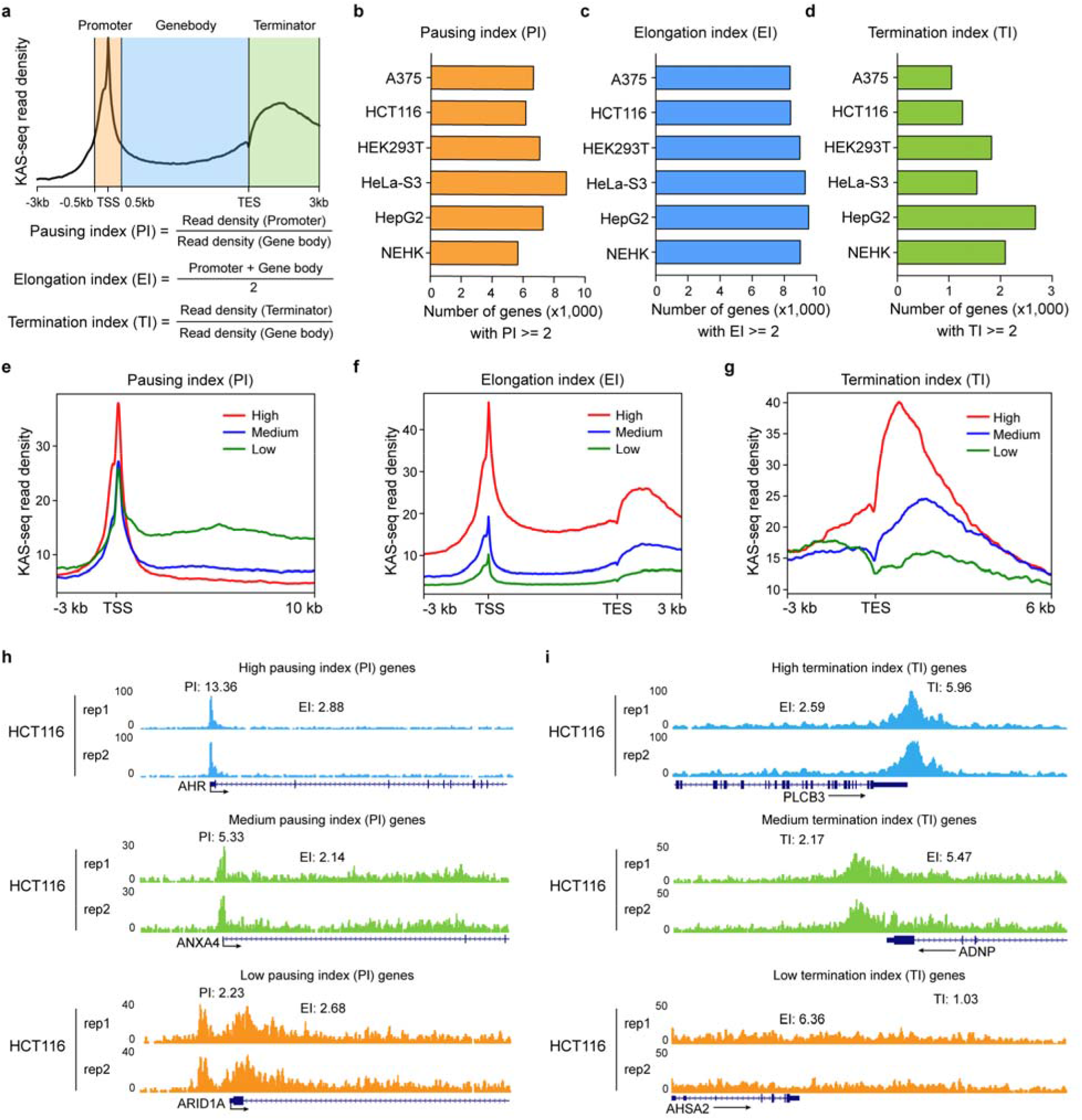
Quantify the dynamics of transcriptionally engaged RNA Pol II. **a** Schematic representation of pausing index (PI), elongation index (EI) and termination index (TI) calculation strategy using mapped KAS-seq reads on gene coding regions. **b-d** Bar plots showing the number of protein-coding genes with PI (b), EI (c) and TI (d) values greater than or equal to 2. PI, EI and TI values were calculated using the KAS-seq data generated in 6 human cell lines. **e-g** Metagene profiles showing the averaged KAS-seq read density on different groups of genes defined by PI (e), EI (f), and TI (g) metrics. Red lines represent genes with high PI, EI and TI; Blue lines represents genes with medium PI, EI and TI; Green lines represents genes with low PI, EI and TI. **h-i** Snapshots of KAS-seq custom tracks in HCT116 cells from UCSC genome browser showing the pattern of KAS-seq reads on the promoters and transcription termination regions of genes with different levels of pausing index (e) and termination index (f). The averaged PI, TI and EI values between two replicates of KAS-seq data were labeled. rep1, replicate1; rep2, replicate2. The arrows represent the transcription direction of the hg19 Refseq annotated genes.

### Estimating the termination length of transcriptionally engaged RNA Pol II

Termination is the final stage of the transcription cycle when RNA Pols and nascent RNA are released from the DNA template, the detailed mechanisms of which are still poorly understood[27]. Improper transcription termination may result in severe consequences; for example, in renal cell carcinoma, transcription read-through results in the aberrant expression of neighboring genes and the generation of chimeric transcripts[28-30]. While the termination index (TI), defined above captures the “releasing” efficiency of RNA Pol II from the DNA template, we also wanted a metric that captures the “braking” efficiency of RNA Pol II over the 3’ end of a protein-coding gene. We therefore used termination length (TL), defined as the width of the KAS-seq peak located in the transcription termination region, to further quantify the efficiency of transcription termination. TI and TL reflect two distinct aspects of the transcription termination process.

TL cannot be precisely captured using current peak-calling methods, as KAS-seq peaks are typically broad (∼2 kb to ∼30 kb). KAS-seq peaks that cover the transcription end site can span two or more genomic features in a gene body or even multiple genes. We propose a strategy to narrow down the location of KAS-seq signals in the transcription termination region. First, KAS-seq peaks are binned into 100 bp sliding windows with 50 bp overlaps. Secondly, sliding windows located in coding regions (spanning from 2 kb upstream of TSSs to TESs) are removed and the remaining sequences are merged to reveal KAS-seq peaks on transcription terminators. Finally, TL is calculated as the length from the TES to the distal-most boundary of transcription-terminator-associated KAS-seq peak. In our benchmark dataset, we observed that the TLs can vary widely across different genes (∼1 kb to ∼20 kb) (Fig. 4a). TLs of most protein-coding genes (57.4% to 65.2%) are between 1 kb and 5 kb (Fig. 4b) and are generally conserved in different cell lines (Fig. 4c-d). In addition, across the transcriptome, we found that TL was not significantly correlated with TI (Pearson correlation coefficients: 0.054 to 0.174) or EI (Pearson correlation coefficients: −0.084 to 0.008) in any of the 6 human cell lines (Fig. 4e-f), indicating that the “releasing” of RNA Pol II from DNA and the “braking” of RNA Pol II’s elongation at terminators may be separate processes that involve different molecular mechanisms.

**Figure 4.**
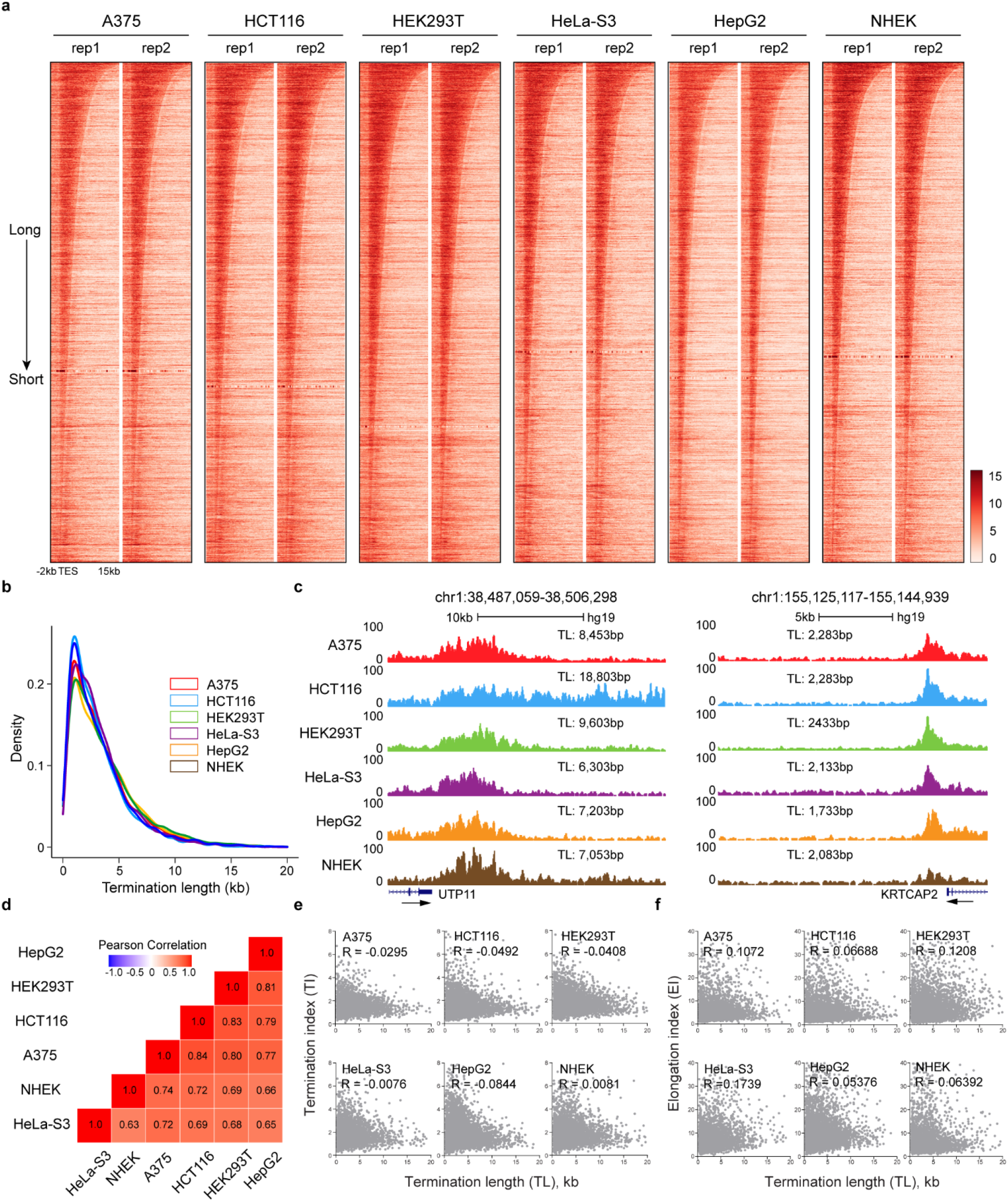
Calculate the termination length of transcriptionally engaged RNA Pol II. **a** Heatmap plots showing the KAS-seq read density at transcriptional termination regions of protein coding genes, with 2□kb upstream of TES and 15□kb downstream of TES shown. Transcriptional termination regions were ranked using the transcription length (TL) calculated using KAS-pipe2. **b** Density curve plot showing the distribution of calculated termination length using KAS-seq data generated in 6 human cell lines. **c** Snapshots of KAS-seq custom tracks from UCSC genome browser showing the pattern of KAS-seq peaks on the transcriptional termination regions of two example genes with long and short calculated termination length. The averaged TL values between two replicates of KAS-seq data were labeled. TL, termination length. **d** Pearson correlation heatmap of TL calculated on common transcription terminators in A375, HCT116, HEK293T, HeLa-S3, HepG2, and NHEK cells. Pairwise pearson correlation coefficients are noted in each square. **e-f** Scatterplots showing the pearson correlation between TL and TI(e) or elongation index (EI) (f) in A375, HCT116, HEK293T, HeLa-S3, HepG2, and NHEK cells, respectively. R values represent Pearson correlation coefficients.

### The identification of single-stranded transcribing enhancers

Transcribed enhancers are short regulatory elements that synthesize enhancer RNAs (eRNAs) by proximal-paused RNA Pol II[31]. These enhancers are generally associated with specific transcription factor binding and experience more long-range interactions than regular enhancers[11, 32]. Numerous studies have reported that transcribed enhancers are more likely to be functional than those identified using only histone marks or chromatin accessibility data. KAS-seq can be used to define a group of transcribed enhancers using ssDNA signals. In previous studies, we defined transcribed enhancers by directly overlapping KAS-seq peaks with enhancers indicated by distal H3K27ac or DNase I hypersensitive sites (DHSs), termed as single-stranded transcribing enhancers. However, this method may produce false positives, since it cannot distinguish KAS-seq peaks mediated by proximal-paused RNA Pol II from those mediated by transcription elongation, both of which can display ssDNA signals on distal H3K27ac peaks and DHSs. To address this issue, KAS-pipe2 introduces a new filtering strategy to select enhancers with proximal-paused RNA Pols by comparing all KAS-seq peaks to nearby regions. KAS-seq peaks mediated by transcription elongation will not be significantly different from the nearby regions and can be eliminated.

For each enhancer that overlaps a KAS-seq peak, we first define two regions of the same size adjacent to the upstream and downstream boundaries of the enhancer as “enhancer shores”. We then calculate the ssDNA read density in the enhancer and its corresponding shores. Enhancers with enriched ssDNA density (1.5-fold by default) compared to their enhancer shores are identified as single-stranded transcribing enhancers. To illustrate this functionality, we applied KAS-pipe2 to the benchmark KAS-seq dataset. As a result, we identified 2,234 single-stranded transcribing enhancers from 26,719 active enhancers (distal H3K27ac peaks) in HepG2 cells, which strongly overlapped with open chromatin regions defined by ATAC-seq (90.8%, 2029/2234) (Fig. 5a-b). This finding is supported by the results from a nascent RNA-based orthogonal assay, NET-CAGE, in which 65.33% (765/1,171) of transcribed enhancers intersect with those by KAS-seq (Fig. 5c). Moreover, KAS-seq identified approximately twice as many transcribed enhancers (2,234 vs 1,171) as NET-CAGE, which is expected, since nascent eRNAs are readily degraded by the nuclear exosome complex soon after their synthesis[33], whereas ssDNAs are stable (Fig. 5d).

**Figure 5.**
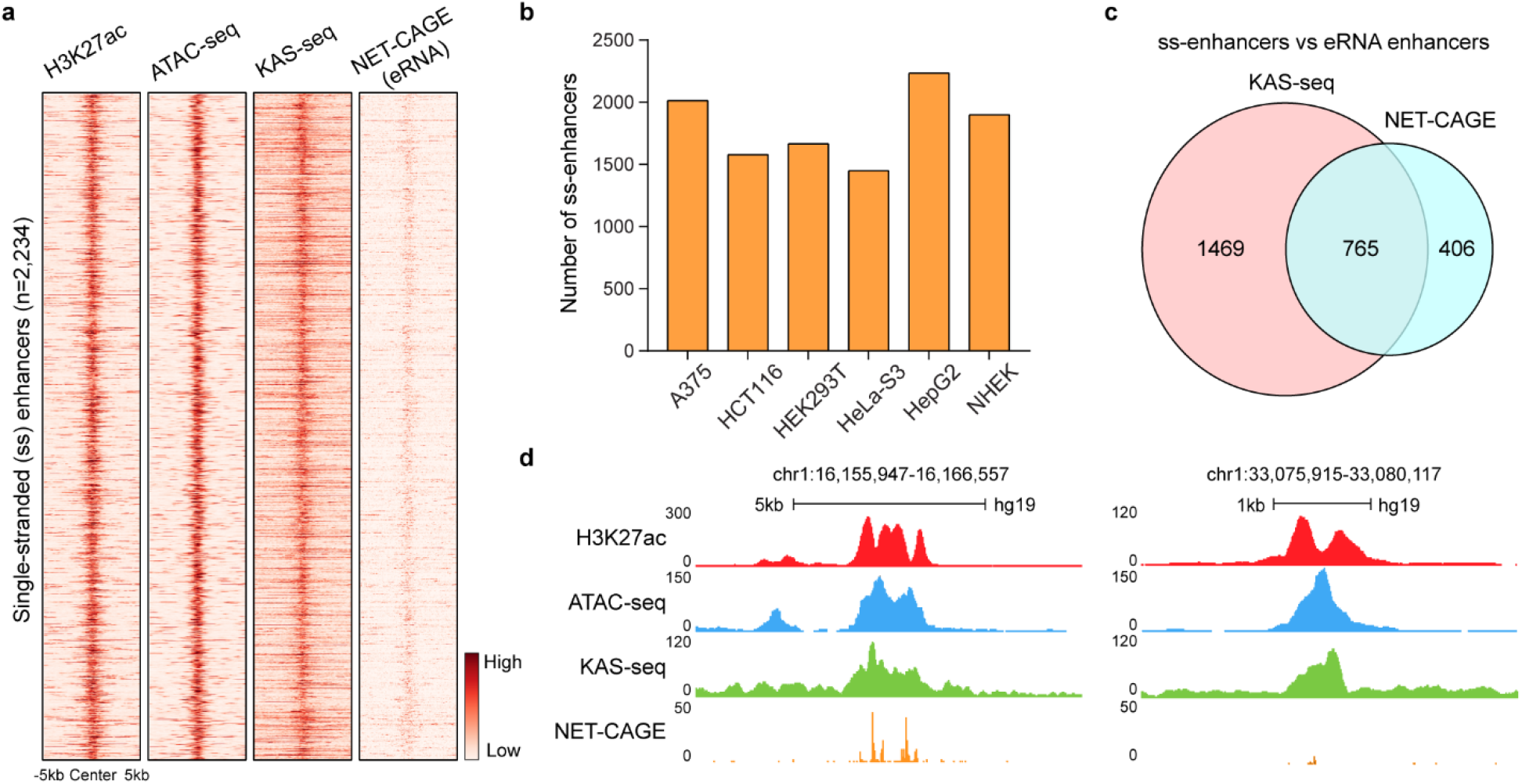
Identify single-stranded transcribing enhancers using KAS-seq data. **a** The heatmap plot showing the KAS-seq and NET-CAGE read density in 2,234 transcribing single-stranded enhancers identified using KAS-seq data in HepG2 cells. Active enhancers were defined as distal H3K27ac peaks identified using ChIP-seq data. **b** Bar plot showing the number of single-stranded transcribing enhancers identified in the benchmark KAS-seq dataset generated in 6 human cell lines. A375: 2,013; HCT116: 1,578; HEK293T: 1,665; HeLa-S3: 1,448; HepG2: 2,234; NHEK: 1,900. **c** Venn diagram showing the common transcribed enhancers defined by KAS-seq and NET-CAGE data in HepG2 cells. **d** Snapshots of custom tracks in UCSC genome browser showing the pattern of H3K27ac, ATAC-seq, KAS-seq and NEG-CAGE peaks on two example single-stranded transcribing enhancers identified using KAS-seq data in HepG2 cells.

### Genome-wide R-loop identification using spKAS-seq data

R-loops are three-stranded structures that consist of an RNA-DNA hybrid and a displaced single-stranded DNA[34, 35]. Functional R-loops form frequently during transcription and play key roles in various nuclear processes, most notably telomere maintenance, DNA repair, transcription regulation, and DNA replication[36]. Previously, genome-wide R-loop detection primarily relied on technologies that enrich for RNA-DNA hybrids using the S9.6 monoclonal antibody or catalytically inactive RNase H before performing high-throughput sequencing, such as DRIP-seq, R-ChIP and mapR[37-40]. spKAS-seq is a strand-specific variant of KAS-seq, developed to map genome-wide R-loop structures in vivo by detecting asymmetric ssDNA exposure on the two strands (Fig. 6a). To map R-loops using spKAS-seq data, KAS-pipe2 requires at least three biological replicates. First, it splits the uniquely mapped reads based on their strand information. Then a read count matrix is constructed for each 500bp sliding window (with 50% overlap) that interacts with spKAS-seq peak using uniquely mapped reads on two strands. For each window, we applied the Wald test from *DESeq2* to detect strand imbalance and obtained p-values adjusted for multiple comparisons. Finally, we merged concatenating windows with strand imbalance of read density to obtain a catalog of R-loops. In our benchmark dataset, only the libraries from HeLa-S3 cells were constructed using the spKAS-seq protocol. We thus applied KAS-pipe2 to the spKAS-seq data in HeLa-S3 cells for R-loop identification. As a result, we obtained 28,578 and 27,033 R-loops on positive and negative strands (Fig. 6c-d), respectively, with most R-loops less than 2.5 kb in length (Fig. 7e). In comparison to results obtained from DRIP-seq, which uses a S9.6 monoclonal antibody, spKAS-seq captures approximately twice as many R-loops (55,611 vs 26,334), a fraction of which were supported by DRIP-seq (26%, 14,438/55,611) (Fig. 6f-g). The discrepancy is probably caused by the fact that DRIP-seq inadvertently perturbs R-loops and is biased toward certain chromatin regions. In summary, our results show that spKAS-seq can be used to generate a comprehensive map of genome-wide R-loops with high resolution and sensitivity[38-42].

**Figure 6.**
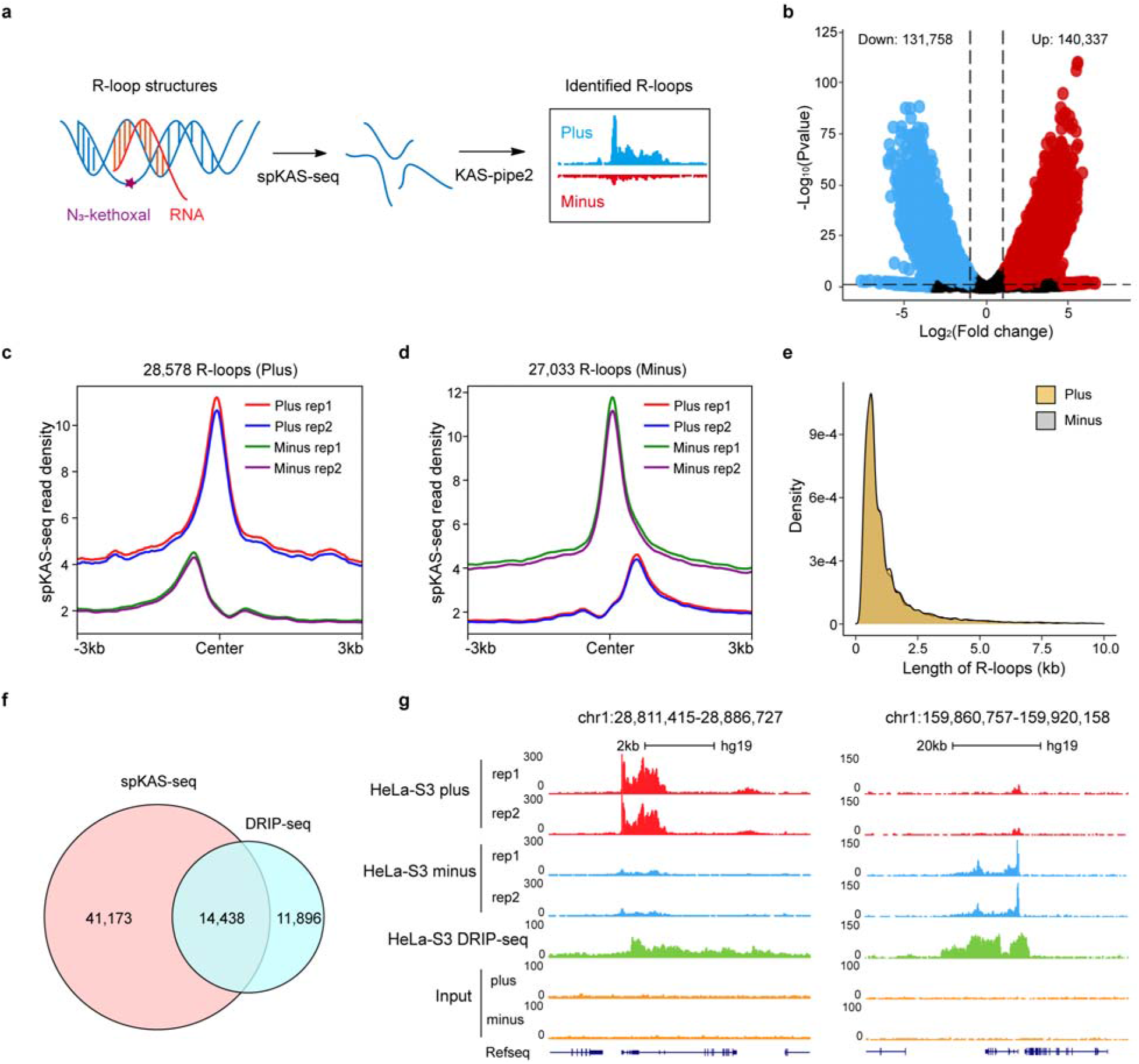
R-loop identification using spKAS-seq data. **a** The schematic diagram spKAS-seq to map the genome-wide R-loop structures. **b** Volcano plot showing the number of sliding windows with unbalanced spKAS-seq reads number mapped on plus or minus strands in HeLa-S3 cells. Red dots indicate the sliding windows with significantly higher spKAS-seq reads number on plus strand (n=140,337). Blue dots indicate the sliding windows with a significantly higher number of spKAS-seq reads on the minus strand (n = 131,758). The sliding windows with unbalanced spKAS-seq read numbers were identified with fold change greater than or equal to 2, and adjusted P value smaller than or equal to 0.05. **c-d** Metagene profile plots showing the spKAS-seq read density with strand specificity on the identified R-loops enriched on plus (b) and minus (c) strands. **e** Density curve plot showing the length distribution of R-loops identified using spKAS-seq in HeLa-S3 cells. Plus: R-loops on plus DNA strand. Minus: R-loops on minus DNA strand. **f** Venn diagram showing the overlapped R-loops identified by spKAS-seq and DRIP-seq data in HeLa-S3 cells. **g** Snapshots of spKAS-seq data custom tracks from UCSC genome browser showing the pattern of unbalanced spKAS-seq reads mapped to plus or minus strand on two example R-loops identified by KAS-pipe2 using spKAS-seq data in HeLa-S3 cells.

**Figure 7.**
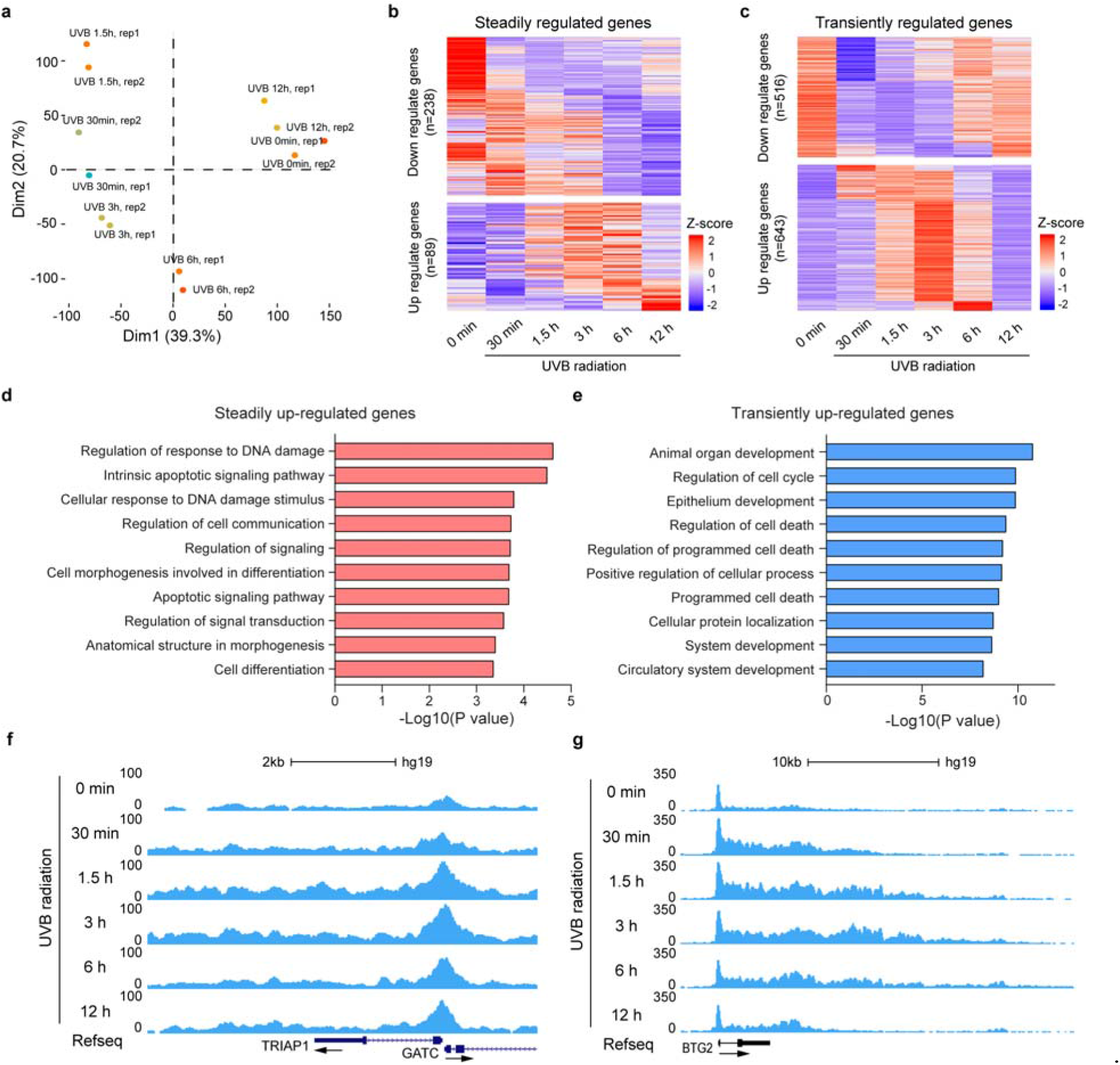
Differential RNA Pol II activity analysis for time-course KAS-seq data in NHEK cells. **a** Principal component analysis (PCA) plot of KAS-seq data generated in NHEK cells treated with UVB radiation for 0□min (sham irradiation), 30□min, 1.5 h, 3 h, 6 h and 12h, respectively. rep1, replicate1; rep2, replicate2. **b-c** Heatmap plots showing the dynamics of global RNA Pol II activity patterns of steadily regulated genes (b) and transiently regulated genes (c) based on normalized read-counts of time-course KAS-seq data. **d-e** Bar plots showing the gene ontology (GO) analysis for steadily up-regulated genes (d) and transiently up-regulated genes (e). GO terms of steadily up-regulated genes are shown as ranked red bars, and GO terms of transiently up-regulated genes are shown as ranked blue bars. **f-g** Snapshots of time-course KAS-seq data custom tracks from UCSC genome browser showing the KAS-seq read density on example steadily (*TRIAP1*) (f) and transiently (*BTG2*) (g) up-regulated genes in NHEK cells treated with UVB radiation at different time points.

### Differential RNA polymerases activity analysis for time-course KAS-seq data

Time-course KAS-seq experiments can provide insights into the regulatory mechanisms of gene expression by capturing transient transcriptional dynamics with temporal resolution[11, 12]. KAS-pipe2 supports differential RNA Pols activity analysis for time-course KAS-seq data. As an example, we applied KAS-pipe2 to a time-course KAS-seq dataset with human epidermal keratinocytes (NHEK) cells treated with UVB radiation for 0□min (sham irradiation), 30□min, 1.5 h, 3 h, 6 h, and 12h, respectively. We observed that UVB radiation treatment clearly induced transcriptional changes from 30□min to 12 h, leading to distinct KAS-seq profiles at each time point (Fig. 7a). From differential time-course RNA Pols activity analysis, we identified 327 genes (89 up and 238 down) as ‘steadily regulated genes’ with continuous effects (Fig. 7b), and 1,161 genes (643 up and 516 down) as ‘transiently regulated genes’ with fleeting effects (Fig. 7c). Moreover, gene ontology (GO) analysis revealed that steadily up-regulated genes were mainly related to DNA damage and apoptosis signaling pathways (Fig. 7d), and transiently up-regulated genes were significantly enriched in critical biological processes, including cell development, cell cycle, and cell death (Fig. 7e). For instance, *TRIAP1* was identified as a steadily up-regulated gene that prevents apoptosis by mediating intramitochondrial transport of phosphatidic acid[43], whereas *BTG2*, which was transiently up-regulated with UVB radiation treatment only for 30 min to 6 h (Fig. 7f-g), is involved in cell cycle processes and functions as an antiproliferative p53–dependent component of the DNA damage cellular response pathway[44].

## Discussion

In this study, we present KAS-pipe2, a user-friendly, one-stop shop workflow to explore KAS-seq and spKAS-seq data. Specifically, we propose several novel metrics to capture the activity of transcriptionally engaged RNA Pol II, including pausing index (PI), elongation index (EI), termination index (TI), and termination length (TL). KAS-pipe2 provides an excellent scheme to study the transient dynamics of each step in the transcription cycle. By examining these metrics in our benchmarking cell lines, we found the termination indexes and termination lengths of same genes are not significantly correlated, indicating that the “releasing” of RNA Pol II from terminators (termination index) and the “braking” of RNA Pol II’s elongation at terminators (termination length) may involve two separate processes. Currently, two models have been proposed for the transcription termination process: (1) RNA Pol II is released following an allosteric change in the elongation complex (the allosteric model); (2) the elongation complex is dismantled following degradation of the nascent RNA transcripts by a 5’–3’ exoribonuclease (the torpedo model). These two models mainly provide mechanistic details of the “releasing” of RNA Pol II from DNA, but not the “braking” of RNA Pol II’s elongation at terminators. We suspect the sequence context may play an important role in the “braking” of RNA Pol II’s elongation, as we found that TLs are generally conserved in different cell lines, which was not observed for TI. In addition, considering the fast (5 min) and specific reaction between N_3_-kethoxal and ssDNA, another possible application of KAS-seq is to quantify the elongation rate of RNA Pol II during transcription through the treatment and release of transcription inhibitors. Besides processing KAS-seq and spKAS-seq data, KAS-pipe2 can also facilitate interpretation of other types of sequencing data. For instance, the QC, read alignment, and differential analysis modules are suitable for ChIP-seq and chromatin accessibility data, and the transcription metrics module is suitable for GRO-seq and PRO-seq data.

Compared to the previous version of the pipeline (KAS-pipe), which only includes a few stand-alone shell scripts[12], KAS-pipe2 adopts a new framework design to implement functionalities under the usage: KAS-pipe2 <subcommand> [options], which is convenient to update with new features and has been adopted in many other popular packages, such as *deeptools* and samtools. KAS-pipe2 not only provides foundational tools for analysis of KAS-seq and spKAS-seq data, but it also includes many expanded features to facilitate the data interpretation. Still, the current workflow does have limitations, especially for the identification of genome-wide R-loops. For instance, non-canonical DNA structures other than R-loops, such as triple-strand DNA (H-DNA), may also asymmetrically expose ssDNA, thereby contributing to a small portion of defined R-loop signals. While the current definition strategy applied in KAS-pipe2 cannot exclude this type of false positive R-loop signal, which can be avoided by using a deconvolution method or by including non-canonical DNA structure annotation. Finally, we plan to enhance future versions of KAS-pipe2 by developing more analytical tools for new applications of KAS-seq experiments. One update we hope to include is a method for the construction of gene regulatory networks using time-course KAS-seq data.

## Conclusion

KAS-pipe2 is a comprehensive tool suite, its application to KAS-seq data enabling researchers to study the dynamic activities of transcriptional engaged RNA Pols and transient transcriptional regulatory programs. Through a benchmark dataset consisting of KAS-seq data generated from 6 human cell lines, we thoroughly showcase novel features implemented in KAS-pipe2 and diverse applications of the KAS-seq technology, including calculation of transcription related metrics, identification of single-stranded transcribing enhancers, identification of R-loops with high resolution, and differential RNA Pols activity analysis. KAS-pipe2 provides a powerful workflow for pre-processing, analysis, and interpretation of KAS-seq and spKAS-seq data.

## Methods

### Cell culture

A375, HCT116 and HepG2 cells were purchased from ATCC (CRL1619 for A375, CCL247 for HCT116, and HB8065 for HepG2) and were cultured in DMEM (Gibco 11995) supplemented with 10% (v/v) fetal bovine serum (Gibco), 1% penicillin and streptomycin (Gibco 10378) and grown at 37 °C with 5% CO2. Epidermal Keratinocytes (NHEK) cells were also purchased from Lonza and maintained in KGM Gold keratinocyte growth basal medium (Lonza, #00192151) and KGM Gold keratinocyte growth medium supplements and growth factors (Lonza, # 00192152). All cell lines were routinely verified to be free of mycoplasma.

### UVB radiation

After washing twice with phosphate-buffered saline (1×PBS buffer, Invitrogen), cells were UVB-irradiated (20 mJ/cm2) using UV Stratalinker 2400 with UVB bulbs (Stratagene). Control samples were treated with sham irradiation. The UVB dose was monitored regularly using the Goldilux UV meter with a UVB detector (Oriel Instruments). Cells were collected at 0□min, 30□min, 1.5 h, 3 h, 6 h and 12h time points for KAS-seq experiments.

### KAS-seq and spKAS-seq experiments

The experimental procedures of N_3_-kethoxal labeling and genomic DNA fragmentation are exactly the same for KAS-seq and spKAS-seq experiments, which have been described previously[11, 12]. In brief, after the completion of gDNA N_3_-kethoxal labeling and fragmentation, 5% of the sonicated DNA was reserved as input and the remaining 95% was used for enrichment with 10 μL Dynabeads MyOne Streptavidin C1 (Thermo, 65001). The beads were washed by 1× B&W buffer (5 mM Tris-HCl pH 7.4, 0.5 mM EDTA, 1 M NaCl, 0.05% Tween-20), resuspended in 95 μL 2× B&W buffer, and mixed with fragmented gDNA. The binding was performed at room temperature for 15 min. The beads were washed once in 1× B&W buffer and then washed twice with 100 mM NaOH solution to denature the dsDNA and remove the DNA strands that were not labeled by N3-kethoxal. (Note: This step is only necessary for spKAS-seq experiments.) A final wash with 1× B&W buffer followed. DNA was eluted in 10 μL H2O by heating the beads at 95 °C for 10 min. Input and enriched DNA were used for library construction by using the Accel-NGS Methyl-seq DNA library kit (Swift, 30024). (Note: The library construction of regular KAS-seq can employ the equivalent alternatives.) KAS-seq or spKAS-seq libraries were then sequenced on the Illumina sequencing platforms. Of the benchmark KAS-seq data in 6 human cell lines, only the HeLa-S3 cell library was constructed in a strand-specific manner.

### Installation of KAS-pipe2

Installation instructions are provided at https://github.com/Ruitulyu/KAS-pipe2. Before applying KAS-pipe2 on the KAS-seq or spKAS-seq data of interest, users first need to download or clone the KAS-pipe2 git repository on github and run ‘bash ./setup.sh’ to make all the scripts executable. Then, users should install all the required software with Conda environments (Unix based platforms) by executing the install tool of KAS-pipe2. Finally, before applying the tools in KAS-pipe2 on real KAS-seq or spKAS-seq data sets, users need to activate the Conda ‘KAS-pipe2’ environment.

### Detailed implementations of main analytical tools in KAS-pipe2

KAS-pipe2 was implemented using BASH scripts, R, and Python. In total, 34 analytical tools in 8 modules were included in the KAS-pipe2 toolkits. All the tools in KAS-pipe2 toolkits can be executed using the command line mode ‘KAS-pipe2 <sub-command> [options]’. The manual of KAS-pipe2 is available at https://ruitulyu.github.io/KAS-pipe2, and detailed descriptions of main analytical tools in KAS-pipe2 are included below.:

### Raw read trimming and alignment

To map KAS-seq or spKAS-seq data confidently to the reference genome of interest, KAS-pipe2 trims off low-quality sequence, adaptor sequence and primer sequence from single-end or paired-end raw FastQ files using the trim_galore package, which can automatically detect the adaptor and primer sequences. In the ‘trim’ tool of KAS-pipe2, 30 bp is by default set as the shortest limit of read length after trimming off the ‘bad’ sequence.

For the read alignment of KAS-seq data, KAS-pipe2 provides two popular and top-ranked aligners, *BWA-MEM* and *Bowtie2*[23, 45], and *Bowtie2* is used as the default aligner. Mapped reads in sam files from the aligners are sorted and converted to bam files using ‘samtools sort’, which are subsequently deduplicated using ‘picard MarkDuplicates’ or ‘samtools rmdup’ (single-end KAS-seq data)[46, 47]. For single-end KAS-seq data, mapped reads were extended to 150bp as default, regardless of the read length of raw sequencing data. For paired-end KAS-seq data, KAS-pipe2 includes a Python script that enables “properly paired” mapped reads to be combined into single interval. An option in the read alignment tools of KAS-pipe2 supports filtering the unique mapped reads with mapping quality ≥10. We also included a ‘spKAS-seq’ tool developed specifically for the read alignment of spKAS-seq data that supports the automatic identification of R-loops without statistical calculations. Details about R-loop identification based on statistical model are described under the “Genome-wide R-loop identification” heading in the Methods section.

### KAS-seq and spKAS-seq peak calling

The “peak calling” tool in KAS-pipe2 calls KAS-seq peaks using the popular and top-performing peak calling software, MACS2[19]. Because KAS-seq data show broad peaks in the gene bodies and transcription termination regions, MACS2 is run by default to call broad peaks by linking nearby enriched regions (--broad) under q-value equal to 0.01. KAS-pipe2 also helps specify the information of effective genome size based on the reference genome for KAS-seq read alignments. To further facilitate confident peak calling, KAS-pipe2 discards KAS-seq reads mapped on problematic regions of the genome in the blacklist.

### Quality control metrics

KAS-pipe2 provides a number of pre-or post-alignment quality control metrics to evaluate KAS-seq data, including library complexity (PCR bottlenecking coefficient and non-redundant fraction), read alignment statistics (numbers of raw reads, clean reads, mapped reads, deduplicated reads and uniquely mapped reads), clean reads rates, alignment rates, duplication rates, fragment size distribution, number of peaks, Pearson correlation coefficients (between replicates or across samples), and the fraction of reads in peaks (FRiP). Clean reads represent raw reads after trimming low-quality and adapter sequence. KAS-pipe2 also supports other types of quality control and generates quality assessment plots, such as saturation analysis, fingerprint plots, genomic distribution of KAS-seq peaks, metagene profiles, and heatmap plots. In KAS-pipe2, the metagene profiles and heatmap plots are generated using the *deeptools* package[20]. In general, high-quality KAS-seq data should have FRiP values greater than 25%, more than 50,000 KAS-seq peaks, and strong KAS-seq peaks in the promoter-proximal and transcription-termination regions.

### Normalization for KAS-seq and spKAS-seq data

The default normalization method used by KAS-pipe2 calculates the reads per kilobase per million mapped reads (RPKM) based on sequencing depth[35]. However, users can customize the analysis by providing scaling factors based on the number of spikeIn reads or other alternative approaches. In KAS-pipe2, we also included a dedicated tool for KAS-seq data normalization, which generates the normalized density files (bedGraph and bigWig).

### Genome annotation

In KAS-pipe2, many tools employ gene annotation for data analysis, such as pausing index, elongation index, termination index and termination length calculation, genomic distribution of spKAS-seq peaks, and enhancer target genes definition. KAS-pipe2 provides the NCBI Refseq gene annotations for many species that are commonly used in real research, e.g. Human: hg18, hg19, hg38; Mouse: mm9, mm10, mm39; C.elegans: ce10, ce11; D.melanogaster: dm3, dm6; Rat: rn6, rn7; Zebra fish: danRer10, danRer11.

### Generation of UCSC genome browser track files

KAS-pipe2 outputs normalized bigWig and bedGraph files for KAS-seq data visualization in a genome browser, such as UCSC and IGV[48, 49]. The “UCSC” tool of KAS-pipe2 generates the bedGraph read density file with track definition lines that can be used to upload and display as custom tracks in the UCSC genome browser.

### Pausing index, elongation index, and termination index calculation

As with GRO-seq[6], we calculate the pausing index (PI) as the ratio of KAS-seq read density in promoter-proximal regions to the density in the gene bodies. Promoter-proximal regions were defined as 0.5 kb upstream and downstream of the transcription start site (TSS), and gene bodies were defined as 0.5 kb downstream of TSS to transcription end site (TES). The ‘bamCoverage’ tool of *deeptools* is used to calculate KAS-seq read density as the number of reads per bin (50 bp), which is then normalized using RPKM[20]. The elongation index (EI) is calculated as the averaged KAS-seq density in the promoter-proximal and gene-body regions. The EI quantitatively measures the activity of transcribing RNA Pols. Of note, promoter-proximal and gene-body KAS-seq density are calculated independently. Similar to the PI, the termination index (TI) is calculated as the ratio of KAS-seq read density at transcription-termination regions to that in the gene bodies. Transcription-termination regions were defined as 3 kb downstream of TES. The ‘index’ and ‘KASindex’ tool in KAS-pipe2 takes the bam files of KAS-seq or spKAS-seq data as input and calculates the KAS-seq read density in promoter-proximal regions, gene bodies, and transcription-termination regions, which are then used to calculate PI, TI or EI values for all the expressed genes with KAS-seq peaks.

### Termination length calculation

Termination length is defined as the width of the KAS-seq peak in the transcription-termination region, and is used to quantify the efficiency of RNA Pols release from their DNA template. First, reference genome is split into 100 bp (as default) sliding windows with 50 bp overlap using Bedtools[50], and KAS-seq peaks overlapped with transcription end site (TES) are used to filter sliding windows. Next, filtered sliding windows located in the gene coding regions (promoters and gene bodies) were discarded, and the remaining sliding windows were then merged to simulate the transcription-terminator-associated KAS-seq peaks. Finally, termination length is calculated as the length from the TES to the distal-most boundary of transcription-terminator-associated KAS-seq peak. The ‘termination length’ tool in KAS-pipe2 takes the KAS-seq peaks and bam files as input and can be executed using the command: KAS-pipe2 termilength -o KAS-seq_termination_length -t 10 -g mm10 -p peaks.txt -l labels.txt -k KAS-seq.txt &

### Single-stranded transcribing enhancers identification

Single-stranded transcribing enhancers broadly refer to active enhancers that overlap with KAS-seq peaks mediated by proximal-paused RNA Pol II. First, to identify single-stranded transcribing enhancers using KAS-seq data, we defined enhancer shores as two same-size regions adjacent to the upstream and downstream boundaries of the enhancer. Next, we calculated the ssDNA read density in each enhancer and its corresponding enhancer shores, using the ‘multiBigwigSummary’ of the *deeptools* package[20]. Finally, enhancers with enriched ssDNA density (1.5-fold by default), as compared to their enhancer shores, were selected as single-stranded transcribing enhancers. In KAS-pipe2, the ‘SST_enhancer’ tool takes KAS-seq bam files and a list of pre-defined active enhancers as inputs and automatically outputs the identified single-stranded transcribing enhancers. The list of pre-defined active enhancers can be defined by distal H3K27ac peaks and DNase I hypersensitive sites (DHSs) or annotated by the ENCODE and Roadmap epigenomics projects[51]. Moreover, KAS-pipe2 includes a ‘motif’ tool to streamline the procedure of transcription factor motif enrichment analysis on single-stranded transcribing enhancers using HOMER software (http://homer.ucsd.edu/homer/motif/)[52].

### Differential RNA Pols activity analysis for time-course KAS-seq data

The ‘TC’ tool in KAS-pipe2 uses bam files of KAS-seq data at different time points as input to generate the read-count matrix. KAS-seq read count can be calculated on promoters, gene bodies, genes, KAS-seq peaks or custom genomic bins with the ‘multiBigwigSummary’ tool of *deeptools* package[20]. The KAS-seq read-count matrix is then automatically passed to ImpulseDE2 package to execute the time-course KAS-seq analysis using a negative binomial noise model with dispersion trend smoothing by DESeq2[21, 22]. Finally, KAS-pipe2 creates lists of ‘steadily regulated genes’ and ‘transiently regulated genes’, and a global heatmap plot used to visualize the KAS-seq read density trajectories at different time points. When using this tool, the user needs to specify the type of differential time-course RNA Pols activity analysis, such as “case_only” or “case_control”.

### Genome-wide R-loop identification

R-loops can be identified genome-wide using spKAS-seq data by detecting the genomic regions with imbalanced read count mapped to two DNA strands, which doesn’t apply on regular KAS-seq data. First, the ‘R-loop’ tool in KAS-pipe2 splits the spKAS-seq mapped reads into two groups, plus- and minus-strand mapped reads. spKAS-seq read-count matrix is then generated on the 500bp sliding windows (default, but can be set to an optional length) with 250bp overlap located in the spKAS-seq peaks. KAS-pipe2 applied the negative binomial generalized linear model derived from DESeq2 to identify the sliding windows with significantly imbalanced reads of two DNA strands using the adjust p-values (q values) smaller than or equal to 0.05[21]. The identified sliding windows were subsequently merged into combined genomic regions for R-loops definition. In addition, the ‘R-loop’ tool also generates the R-loop density file by calculating the difference of spKAS-seq read density between two DNA strands on 50 bp genomic bins.

## Declarations

### Ethics approval and consent to participate

Not applicable

### Availability of data and materials

KAS-pipe2 was released under the MIT License and is available at https://github.com/Ruitulyu/KAS-pipe2. A copy of the source code has also been deposited in Zenodo (DOI: https://doi.org/10.5281/zenodo.6519166). Raw sequencing data of KAS-seq and spKAS-seq experiments in benchmark A375, HCT116, HepG2, HeLa-S3 and NHEK cells have been deposited at Gene Expression Omnibus (GEO) under the accession number: GSE202044. Published KAS-seq data in HEK293T cells used in this study can be downloaded from GEO under the accession number: GSE139420.

### Authors’ contributions

R.L. and C.H. conceived and designed the study. R.L. designed and implemented the KAS-pipe2 software on KAS-seq data. T.W. performed the experiments. G.P. and Y.H. generated KAS-seq data in NHEK cells treated with UVB radiation. R.L. and M.C. wrote the paper with the input from all authors. M.C. supervised the overall study. All authors read and approved the final manuscript. The University of Chicago has filed a patent application for KAS-seq with Tong Wu, Ruitu Lyu and Chuan He as inventors.

## Acknowledgements

We thank Shun Liu for thoughtful discussion about the framework design of KAS-pipe2, and other members in Chuan He’s group for feedback on this study. We thank Dr. John M. Gaspar to provide the python script ‘SAMtoBED’, which combines “properly paired” alignments into a single BED interval.

